# Increasing motor skill acquisition by driving theta-gamma coupling

**DOI:** 10.1101/2019.12.20.883926

**Authors:** Haya Akkad, Joshua Dupont-Hadwen, Amba Frese, Irena Tetkovic, Liam Barrett, Sven Bestmann, Charlotte J Stagg

## Abstract

Skill learning is a fundamental adaptive process, but the mechanisms remain poorly understood. Hippocampal learning is closely associated with gamma activity, which is amplitude-modulated by the phase of underlying theta activity. Whether such nested activity patterns also underpin skill acquisition in non-hippocampal tasks is unknown. Here we addressed this question by using transcranial alternating current stimulation (tACS) over sensorimotor cortex to modulate theta-gamma activity during motor skill acquisition, as an exemplar of a non-hippocampal-dependent task. We demonstrated, and then replicated, a significant improvement in skill acquisition with theta-gamma tACS, which outlasted the stimulation by an hour. Our results suggest that theta-gamma activity may be a common mechanism for learning across the brain and provides a putative novel intervention for optimising functional improvements in response to training or therapy.

## Introduction

The acquisition of motor skills is a central part of our everyday lives, from learning new behaviours such as riding a bike, to the recovery of function after brain injury such as a stroke^1–4^. Better understanding of the mechanisms underpinning skill acquisition, and developing mechanistically-informed strategies and tools to promote skill learning in healthy and pathological movement, is therefore a high priority scientific and clinical goal.

Acquisition of motor skills is linked to a number of cortical and subcortical brain regions, but among these, primary motor cortex (M1) is thought to play a central role^1,2,4,5^, making this a key target for neurorehabilitative interventions^6–8^. However, the neurophysiological changes supporting skill acquisition in M1 are poorly understood, substantially hampering the development of novel interventions.

Outside the motor domain, the mechanisms underpinning learning have been most extensively studied in the hippocampus, where theta-amplitude-coupled mid-gamma frequency activity (θ-γ phase-amplitude coupling; PAC) has been hypothesised as a key learning-related mechanism. In rodent hippocampal area CA1, oscillations in the θ (5-12 Hz) band become dominant during active exploration^9^, and have been widely hypothesised to allow information coming into CA1 from distant regions to be divided into discrete units for processing^10,11^. A prominent feature of hippocampal theta activity is its co-incidence with higher-frequency activity in the γ range (30-140 Hz). Gamma coherence in the hippocampus alters during learning^12^ and memory retrieval^13^, and its relative synchrony during task predicts subsequent recall^14,15^.

Hippocampal activity at different frequencies within the gamma band is also coupled to different, specific phases of the underlying theta rhythm, suggesting that the precise relationship between gamma activity and theta phase may be important for function^16–18^. For example, 60-80 Hz activity, which increases significantly during memory encoding, is coupled to the peak of the underlying theta oscillation^19^.

θ-γ PAC appears to be a conserved phenomenon across the cortex, and has been hypothesised as a fundamental operation of cortical computation in neocortical areas^20^. For example, in the sensory cortices, it provides a neural correlate for perceptual binding^21^. In the motor cortex, gamma oscillations at approximately 75 Hz are observed during movement^22–28^, and an increased 75 Hz activity has been observed in dyskinesia, suggesting a direct pro-kinetic role^29,30^. As in the hippocampus, M1 gamma oscillations are modulated by theta activity, with 75 Hz activity in human M1 being phase-locked to the peak of the theta waveform^31^.

However, whether theta-gamma coupling plays a similar role in learning in neocortical regions as it does in the hippocampus has not yet been determined. We therefore wished to test the hypothesis that θ-γ PAC is a conserved mechanism for learning across the brain. To investigate the functional role of θ-γ PAC in learning outside the hippocampus, we modulated local theta-gamma activity via externally applied alternating current stimulation (tACS) over M1 during learning of an M1-dependent skill^32^, which shows robust behavioural improvements over a relatively short period of time.

We reasoned that if θ-γ PAC is a key mechanism for motor skill learning, then interacting with θ-γ PAC, specifically with 75 Hz gamma activity applied at the theta peak^19,31^, via tACS, should have the capacity to delay or speed-up skill acquisition in healthy human participants. Moreover, if the functional role of this theta-gamma PAC is indeed critically dependent on the gamma activity occurring at a specific phase of theta activity, then any behavioural effect should be specific to the theta phase at which the gamma was applied. To address this question, we therefore derived a waveform with gamma applied during the trough of the theta activity as an active control.

## Results

104 healthy participants performed a M1-dependent ballistic thumb abduction task with their left hand^32–34^ while tACS was applied over the right M1. Volunteers trained to increase thumb abduction acceleration in their left, non-dominant, thumb over 5 blocks of 70 trials each (figure 1). We first conducted an exploratory single-blinded experiment, in which we tested for the influence of theta-gamma coupled stimulation on skill acquisition. This experiment revealed that when applied externally over right M1, gamma coupled to the peak of a theta envelope (TGP) substantially enhanced motor skill acquisition, compared to sham and an active stimulation control. Based on these results, we conducted a second, double-blind, pre-registered, sham-controlled experiment, which confirmed the beneficial effect of TGP on motor skill acquisition.

**Figure 1:**
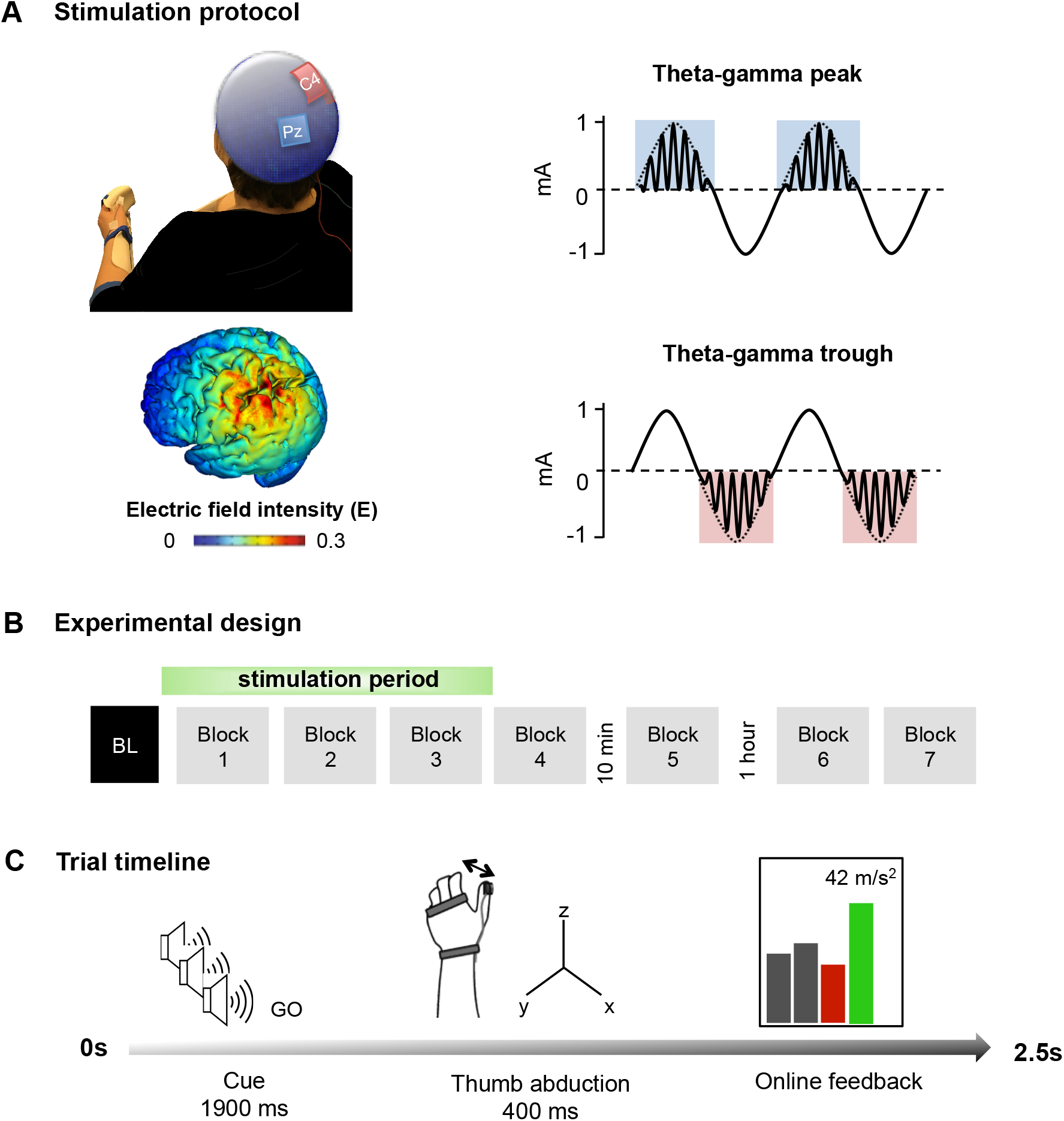
Theta-gamma tACS protocol and task. A. The theta-gamma tACS montage was delivered over right M1, with the anode over right M1 (red, C4) and the cathode over the parietal vertex (blue, Pz). Electrical field distribution projected on a rendered reconstruction of the cortical surface in a single individual. A 75 Hz gamma rhythm was amplitude-modulated by the peak (TGP) or trough (TGT) envelope of a 2mA peak-to-peak 6 Hz theta rhythm. **B.** The task involved 5 blocks, excluding baseline (BL), in experiment 1, and 7 blocks in experiment 2. Each block consisted of 70 trials and inter-block intervals lasted 2 minutes, apart from a 10 minute and 1 hour break after blocks 4 and 5 respectively. Stimulation was delivered for 20 minutes during the first 3. **C.** All trials began with three auditory warning tones acting as a ready-steady-go cue. At the third tone, participants abducted their thumb along the x-axis as quickly as possible and were given online visual feedback of their performance via a screen positioned in front of them. Feedback was presented as a scrolling bar chart with the magnitude of acceleration displayed on a trial-by-trial basis; a green bar indicated acceleration was higher than the previous trial and a red bar indicated the opposite. BL: baseline.

### Experiment 1

58 participants (age: 24 ±5.1 years, 37 female) participated in Experiment 1, and were randomly assigned to one of three experimental groups, which received either 20 minutes of tACS over right primary motor cortex, or sham. For the active tACS condition, participants received (1) theta-gamma peak stimulation (TGP; figure 1A), whereby gamma frequency (75 Hz) stimulation was delivered during the peak of a 6 Hz theta envelope as is found naturally in the human motor cortex^31^, or (2) an active control, theta-gamma trough (TGT) stimulation, whereby the gamma stimulation was delivered in the negative half of the theta envelope. For sham stimulation, 6 Hz theta was briefly ramped up for 10 s, and then ramped down again. Participants performed the skill learning task during the stimulation, and for approximately 15 min after cessation of stimulation.

#### Theta-gamma-peak stimulation significantly improves motor performance

We first wished to assess whether participants were able to improve their performance across the task, regardless of stimulation. As expected, skill increased in all three groups over the course of the experiment [Repeated Measures ANOVA with one factor of Block (1-6) and one factor of Stimulation (TGP, TGT, Sham), Main Effect of Block F(2.203,121.187) = 85.122, p<0.001]. However, the stimulation groups differed significantly in their skill acquisition [Main Effect of Condition F(2, 55)=3.396, p=0.41; Condition*Block Interaction (F(4.407, 121.187)=158.225, p=0.03; Simple Effect of Condition during stimulation F(2,55)=4.13, p=0.021). In line with our primary hypothesis, follow-up analyses demonstrated a 26% larger acceleration gain from baseline during TGP stimulation, compared with sham condition (Independent t-test t(36)=3.052, p=0.004, Cohen’s d=0.98, figure 2A). There was no significant difference between the TGT and sham groups (t(36)=-0.838, p=0.408).

**Figure 2:**
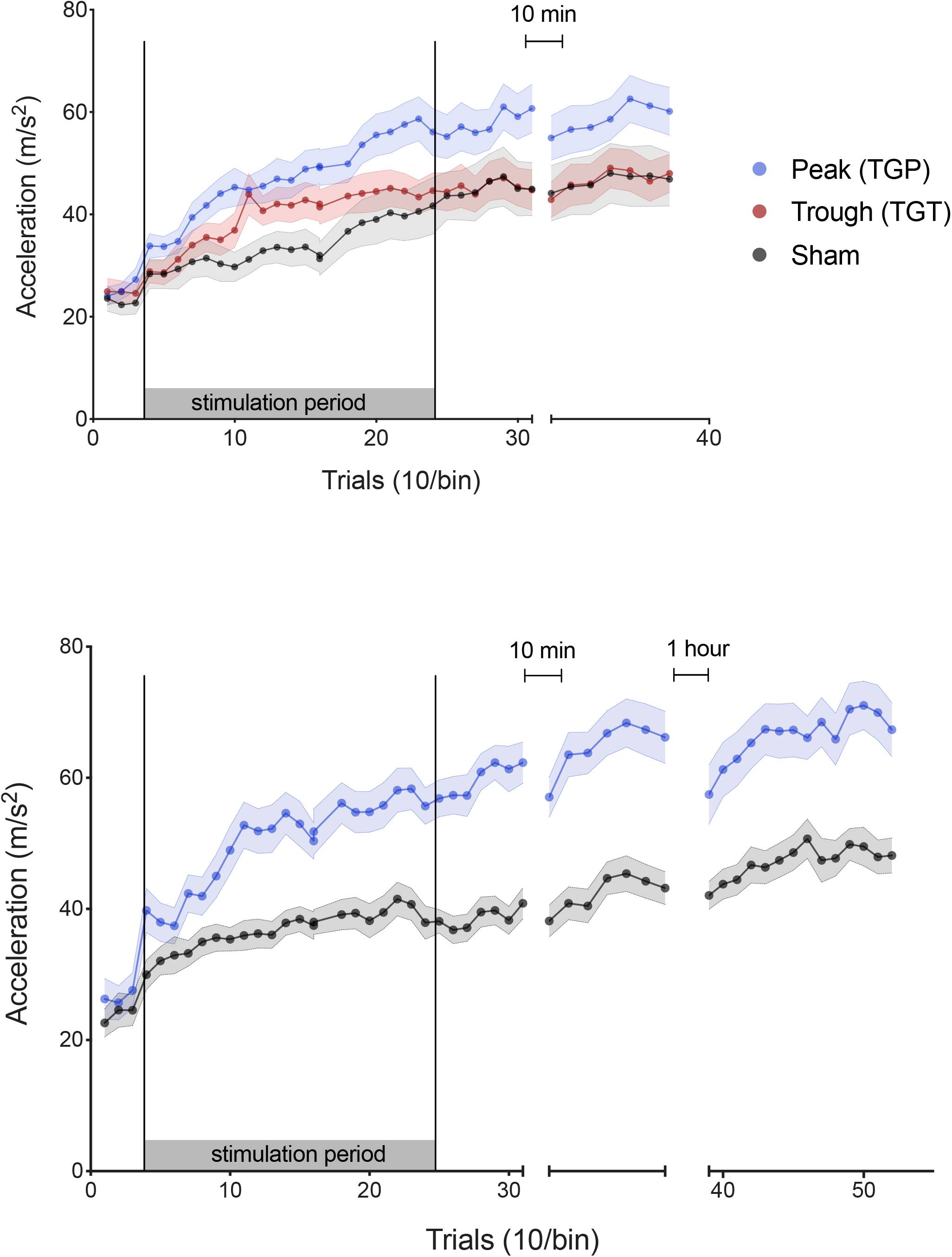
TGP-tACS enhances motor skill acquisition. Mean ballistic thumb abduction acceleration for each stimulation condition. Each point represents the mean of 10 trials across participants and the error bars depict the standard error between participants. **A. Experiment 1.** During stimulation, TGP significantly increased learning over the course of the experiment (i.e., acceleration gain), compared to sham and TGT. After stimulation, the TGP group maintained greater acceleration compared to sham and TGT, but this was no longer significant. **B. Experiment 2.** When replicated in an independent sample, TGP stimulation again substantially enhanced acceleration gain, compared to sham, during stimulation. This effect was maintained 1 hour 15 mins post-stimulation.

There were no significant differences in baseline performance between TGP, TGT and Sham conditions [Repeated measures ANOVA with one factor of block (1-6) and one factor of condition (TGP, TGT, Sham), Simple Effect of Condition within baseline Block F(2,55)=0.30, p=0.743].

#### Motor skill gains are retained post-stimulation

We then wished to explore whether the behavioural effects of stimulation out-lasted the stimulation period, or whether learning in this group returned to baseline after stimulation had ceased. 15 min post-stimulation, the TGP group continued to achieve higher accelerations compared to sham [t(36)=1.787, Cohen’s d=0.58] and TGT [t(38)=2.195, Cohen’s d=0.70], but this was not statistically significant (RM ANOVA Simple Effect of Condition within post-stimulation Block F(2,55)=2.43, p=0.097).

This first experiment established the relevant role of theta-gamma coupled tACS over M1 on motor skill learning in healthy participants, here expressed through an increase in learning. This effect was most effective when gamma frequency stimulation was coupled to the peak of the underlying theta frequency stimulation waveform, as opposed to when it was coupled to the trough of theta. We next sought to confirm this result in an independent cohort, and to further assess the duration of this improvement post stimulation.

### Experiment 2

In order to try to replicate our results from experiment 1, we conducted a double-blind, pre-registered (osf.io/452f8/) replication experiment in an independent sample of 46 participants (age 24±4.1, 32 female, all right handed). Because our first experiment had shown the largest effect on skill learning with TGP stimulation, we now focussed on this condition. Participants were randomised to either TGP stimulation or sham. The experimental protocol was identical to Experiment 1, except that we additionally included a probe to test retention at 1 hour after the end of stimulation. There was no significant difference in baseline performance between TGP and Sham conditions (t(44)=0.734, p=0.467).

As in experiment 1, participants in both conditions showed an improvement in performance throughout the experiment [Repeated measures ANOVA, one factor of Block (1-8), one factor of Condition (TGP, sham); Main Effect of Block F(3.302,145.239) = 72.912, p < 0.001; figure 2B].

However, there was a significant difference in skill acquisition between the two conditions (Main Effect of Condition (F(1,44) = 27.241, p<0.001; Block*Condition interaction F(3.302,145.239) = 7.258, p<0.001). The TGP group achieved significantly greater acceleration gain compared to sham during stimulation [t(44) = 4.201, p <0.001, Cohen’s d = 1.24] and at 15 min post stimulation [t(44) = 4.569, p <0.001, Cohen’s d = 1.35] and this remained significant 75 min post-stimulation[t(44) = 3.430, p = 0.001, Cohen’s d = 1.01]

#### Blinding

Bang’s Blinding Index^35^ indicated successful blinding in both real and sham stimulation groups. Blinding indices were 0.07 and −0.03 in the TGP and sham groups respectively.

## Discussion

Theta-amplitude modulated gamma activity may provide an important mechanism for non-hippocampal-dependent skill acquisition. Using non-invasive brain stimulation to modulate θ-γ PAC in human primary motor cortex in two separate cohorts, one a pre-registered, double-blind study, we demonstrated that externally applied θ-γ PAC during a motor task increases skill acquisition in healthy adults. This behavioural improvement was critically-dependent on the phase-relationship of the theta and gamma components of the stimulation.

θ-γ PAC has consistently been demonstrated to relate to learning in the rodent CA1^16–19,^ and a recent human study demonstrated an improvement in memory using tACS^36^. However, the role of θ-γ PAC role in non-hippocampal-dependent learning had yet to be explored.

Physiologically, M1 gamma activity centered around 75 Hz occurs at the peak of ongoing theta activity^31^ and is ubiquitous in studies of human movement. It only occurs during actual, rather than imagined, movement^27^, and shows topographical specificity within M1^26^. Its hypothesised pro-kinetic role is further supported by the finding of a pathological increase in narrow-band 75 Hz activity within M1 in hyperkinetic patients with Parkinson’s Disease^29^.

Decreases in M1 GABAergic activity are a central mechanism for motor plasticity^37–41^. However, it is not yet clear *how* these decreases alter behaviour. θ-γ PAC may be a candidate mechanism for this: M1 gamma activity arises from GABAergic inter-neuronal micro-circuits involving layer V Parvalbumin +ve neurons^42–49^ are thought to be involved in motor learning^50^. In slice preparations, theta-gamma coupling within M1 arises spontaneously from layer V when GABA activity is blocked^51^. In humans, modulating M1 75Hz activity in humans using tACS leads to a decrease in local GABAergic activity, the magnitude of which predicts motor learning ability on a subject-by-subject basis^28^.

Given the extensive evidence for decreases in GABAergic activity for motor cortical plasticity, it may be that gamma activity, particularly synchronisation of gamma activity via theta oscillations, represents an emergent signature of learning that might be targeted to improve behaviour, though of course the cellular and layer-specificity of our findings remains open.

Here, we tested an *a priori* hypothesis about theta-gamma PAC, and its role in non-hippocampal dependent skill acquisition. We did not test other frequency couplings, and so we cannot claim that similar effects would not be seen with other cross-frequency stimulation paradigms. In addition, θ-γ PAC has been suggested as a mechanism by which anatomically-distant brain regions become functionally connected. We deliberately chose a task that is M1-dependent, thereby providing us with a cortical target for our stimulation, and have not set out to target more than one node in the network. However, this does not imply *per se* that our behavioural effects do not result from effects at multiple nodes within a skill-learning network, and indeed may depend on interactions between multiple brain regions. This hypothesis remains to be tested.

There has been some recent controversy about the contribution of direct stimulation of the underlying neural tissue versus other mechanisms^52,53^ to the behavioural and physiological effects of tACS. While this paper does not aim to directly address this question, two reasons support the notion that our behavioural effects result from direct effects of the current in the brain. Firstly, tACS at the current densities used here have been demonstrated to entrain single-neuron activity in non-human primates^54^, suggesting at least that direct neuronal entrainment is a possible mechanism. Secondly, although stimulation of peripheral scalp nerves has recently been suggested as a putative explanation for behavioural effects of tACS^52^, in experiment 1, we used an inverted waveform as an active control to rule out effects driven by peripheral stimulation, and in experiment 2, there was successful blinding to stimulation type. Collectively, this suggests that the sensory sensations that may arise from stimulation did not substantially differ between active and sham conditions.

In conclusion, we wished to test whether theta-gamma PAC was an important mechanism in non-hippocampally dependent learning in humans. Using a novel non-invasive brain stimulation approach in humans that emulates known neurophysiological activity patterns during learning^19,21,31^, we demonstrated, and then replicated, a substantial behavioural improvement due to stimulation. While the neural underpinnings of this functional outcome need to be explored, this result offers a new technique not only to understand physiological mechanisms of human neuroplasticity, but also potentially a putative novel adjunct therapy for promoting post-stroke recovery.

## Methods

104 healthy participants (Experiment 1: 58 participants, 24 years ±5.1, 37 female; Experiment 2: 44 participants, age 24 ±4.1, 32 female) gave written informed consent to participate in the experiment in accordance with local ethics committee approval. Participants were right-handed and had no contraindications for tACS. Participants visited the lab on one occasion, when they were asked to perform a ballistic thumb abduction task (Figure 1C), the aim of which was to increase their maximum thumb acceleration as much as possible.

### Experiment 1

Participants were randomly assigned to one of three tACS conditions (N= 20 per condition): (1) theta-gamma peak stimulation (TGP; figure 1A), whereby gamma frequency (75 Hz) stimulation was delivered during the peak of a 6 Hz theta envelope, (2) an active control: theta-gamma trough (TGT) stimulation where the gamma stimulation was delivered in the negative half of the theta envelope, and (3) sham stimulation. Participants were blinded to the type of stimulation delivered and naïve to the purpose of the experiment.

#### Experimental set-up

Participants performed a ballistic thumb abduction training task requiring abduction of their left (non-dominant) thumb with maximal acceleration^32–34^. Participants were seated with their left arm slightly abducted, with the elbow flexed to 45**°** (where 0**°** is full extension) and the forearm semi-pronated with the palm facing inwards. The left hand was chosen to avoid ceiling effects that might be present in the dominant hand. The arm, wrist and proximal interphalangeal joints were secured in a plastic custom-built arm fixture to prevent the unintentional contribution of whole hand movement to the ballistic acceleration, though the thumb was left free to move (figure 1C).

The acceleration of the thumb was measured in the x-axis (abduction plane) using an accelerometer (ACL300; Biometrics Ltd., UK) attached to the distal phalanx of the thumb. Recording from the accelerometer was confined to one axis to allow for good skill improvement by providing simplified feedback for the participant^32–34^.

#### Behavioural task

Participants performed ballistic thumb abduction movements of their left hand at a rate of 0.4 Hz indicated by a ready-steady-go procedure, with each of three auditory tones (400 Hz, 300 ms duration) spaced at 500 ms intervals. Participants were instructed to move their thumbs at the onset of the third auditory tone. The behavioural task was separated into 6 blocks (figure 1B). Participants performed an initial baseline block of 30 trials. This was followed by 4 blocks separated by a break of at least 2 min to minimize fatigue, and a final block separated by a 10 min break. Each of these five blocks consisted of 70 trials with a 30 s break between every 35 trials to avoid within block fatigue. Participants were asked to remain at rest during breaks, avoiding any thumb movement.

In all blocks except the baseline block, participants were instructed to move as fast as possible and were encouraged to try to increase their acceleration on every trial. Participants were given visual feedback about the acceleration of their movements on a trial-by-trial basis (figure 1C). Feedback was presented as a scrolling bar chart with the magnitude of the current acceleration plotted after each trial. If the acceleration on the current trial was greater than on the previous trial, the bar was plotted in green, and if it was less the bar was plotted in red. If a movement was made too early or too late (i.e. movement outside a 300 ms window centred on one second after the first tone), no acceleration feedback was given, instead, the message “too early” or “too late” respectively was presented. Additionally, participants were informed of their progress by displaying a moving average of acceleration values over the preceding 10 trials, indicated by a line plotted on the screen over the locations of the 10 consequential trials.

In the baseline block, participants were told to move as closely as possible to the onset of the third tone, and feedback about the temporal accuracy of the movement was given by the experimenter.

#### Behavioural data analysis

Data were analysed via Matlab (Mathworks). The maximal acceleration was calculated for each trial, and any trials with a maximum acceleration less than 4.9ms^2^ were rejected^32^. Additionally, if movements were made too early or too late, i.e. the onset of acceleration of the movement lay more than 300 ms before or after the expected movement time, they were also rejected^32^. Together, this approach led to 1.45±0.94 (mean ± SD) trials being removed per block of 70 trials in experiment 1, and 0.88±0.99 (mean ± SD) trials removed per block of 70 trials in experiment 2. There was no statistical difference between the number of trials being removed per block in each condition (Experiment 1: Mixed ANOVA, block*condition (F(5.409, 148.742)=1.649, p=0.145; Experiment 2: Mixed ANOVA, block*condition (F(2.8, 137.4)=1.05, p=0.396).

#### Transcranial alternating current stimulation (tACS)

tACS was delivered via a DC stimulator in AC mode (NeuroConn DC-Stimulator Plus) through a pair of sponge surface electrodes (5 x 5 cm^2^). Saline was used as a conducting medium between the scalp and the electrodes. The anode was centred over the right primary motor cortex (C4) and the cathode was positioned over the parietal vertex (Pz), in accordance with the international 10-20 EEG system. Impedance was kept below 10 kΩ. The electrode positions were based on simulation of current flow across the brain, using HD-Explore™ software (Soterix Medical Inc., New York) which uses a finite-element-method approach to model electrical field intensities throughout the brain^55^. This confirmed that current was directed to the primary motor cortex (figure 1A).

The theta-gamma-peak (TGP) condition consisted of 20 min continuous, sinusoidal 6 Hz (theta) stimulation at an intensity of 2 mA peak-to-peak, coupled with bursts of a sinusoidal 75 Hz (gamma) rhythm amplitude-modulated by the positive theta phase (0-180°; figure 1A). The theta-gamma-trough (TGT) condition consisted of 20 min continuous, sinusoidal 6 Hz (theta) stimulation at an intensity of 2 mA peak-to-peak, coupled with bursts of a sinusoidal 75 Hz (gamma) rhythm amplitude-modulated by the negative theta phase (180°-360°). Finally, the sham condition consisted of a 10 s continuous sinusoidal 6 Hz stimulation.

The theta-gamma waveforms were custom-coded on the Matlab software and delivered to the NeuroConn stimulator via a data acquisition device (National Instruments USB-6259 BNC). Theta-gamma stimulation was then delivered to the scalp surface electrodes through the NeuroConn stimulator in ‘remote’ mode. Sham stimulation was delivered directly through the NeuroConn stimulator.

tACS was administered in a between-subject design. Participants were randomized to receive either 10 s of sham stimulation during the 1^st^ training block or 20 min of TGP or TGT stimulation during the first 3 training blocks. Participants were blinded to the stimulation condition used and naïve to the purpose of the experiment.

#### Statistical analyses

Data were tested for normality using the Kolmogorov-Smirnov test. Statistical analyses were performed using SPSS. We used a two-way mixed ANOVA with 2 independent variables, ‘condition’ (between-subject variable) and ‘block’ (within-subject variable). Acceleration in ms^2^ was our only dependent variable. Where there was a significant Block*Condition interaction, we analysed the Simple Effect of Condition within levels of Block. Post-hoc t-tests were conducted as appropriate and multiple comparisons were corrected for using the Tukey HSD test. When sphericity assumptions were violated, results are reported with a Greenhouse-Geiser correction.

### Experiment 2

We performed a pre-registered, double-blinded replication of experiment 1 (theta-gamma-peak and sham only) in an independent sample. The experimental design was pre-registered in full on the Open Science Framework (osf.io/452f8/). The experimental design was identical to Experiment 1, except for the following aspects:

#### Power calculation

Sample size was calculated based on the Cohen’s d effect size of the mean improvement in performance from baseline between the theta-gamma-peak and sham conditions in Experiment 1. Given a Cohen’s d = 0.98, 1-β = 0.95 and α = 0.05 this gave a sample size of 24 per group (G*Power), and allowing for a 10% loss of data, we recruited 27 participants per condition.

#### Blinding

On the day of testing, a researcher not involved in data analysis and blinded to experimental protocol and rationale (A.F, I.T, L.B) collected the data and interacted with the participant. Another researcher (HA), not involved in data collection and blinded during data analysis, set-up the stimulation condition on the day of testing, but did not interact with the participant. Unblinding was performed following the completion of data collection and analysis. Participants were naïve to the purpose of the experiment.

Participants completed a blinding questionnaire at the end of the experiment that required them to identify whether they believed they had received real or sham stimulation. To assess the effectiveness of our blinding, we used Bang’s blinding index (BI), where a BI of 1 suggests complete unblinding, a BI of 0 random guessing and a BI of −1 opposite guessing.

#### Behavioural task

The behavioural task parameters were identical to those in experiment 1, but now with an additional 2 training blocks separated from the previous 6 blocks by a break of 1 hour (Figure 1B). During the 1 hour break, participants remained seated and at rest while watching a documentary (Planet Earth, season 1 episode 10). The additional 7^th^ and 8^th^ blocks were separated by a break of at least 2 min to minimize fatigue and each consisted of 70 trials, with a 30 s break between every 35 trials to minimize within block fatigue. Participants were asked to remain at rest during breaks, avoiding any thumb movement.

#### Transcranial Alternating Current Stimulation

Stimulation parameters were identical to those in experiment 1, but only included the theta-gamma-peak (TGP) and sham conditions. Both the participant and the experimenter were blinded to the stimulation condition used.

## Acknowledgements

SB was funded by Brain research-UK (201617-03). CJS holds a Sir Henry Dale Fellowship, funded by the Wellcome Trust and the Royal Society (102584/Z/13/Z). The work was supported by the the NIHR Biomedical Research Centre, Oxford and the NIHR Oxford Health Biomedical Research Centre. The Wellcome Centre for Integrative Neuroimaging is supported by core funding from the Wellcome Trust (203139/Z/16/Z).

## Notes

https://www.osf.io/452f8/

